# High-throughput transposon mutagenesis in the family Enterobacteriaceae reveals core essential genes and rapid turnover of essentiality

**DOI:** 10.1101/2022.10.20.512852

**Authors:** Fatemeh A. Ghomi, Gemma C. Langridge, Amy K. Cain, Christine Boinett, Moataz Abd El Ghany, Derek J. Pickard, Robert A. Kingsley, Nicholas R. Thomson, Julian Parkhill, Paul P. Gardner, Lars Barquist

## Abstract

The Enterobacteriaceae are a scientifically and medically important clade of bacteria, containing the gut commensal and model organism *Escherichia coli*, as well as several major human pathogens including multiple serovars of *Salmonella enterica* and *Klebsiella pneumoniae*. Essential gene sets have been determined for several members of the Enterobacteriaceae, with the *E. coli* Keio single-gene deletion library often regarded as a gold standard for gene essentiality studies. However, it remains unclear how gene essentiality varies between related strains and species. To investigate this, we have assembled a collection of thirteen sequenced high-density transposon mutant libraries from five genera within the Enterobacteriaceae. We first benchmark a number of gene essentiality prediction approaches, investigate the effects of transposon density on essentiality prediction, and identify biases in transposon insertion sequencing data. Based on these investigations we develop a new classifier for gene essentiality. Using this new classifier, we define a core essential genome in the Enterobacteriaceae of 201 universally essential genes, and reconstruct an ancestral essential gene set of 296 genes. Despite the presence of a large cohort of variably essential genes, surprisingly we find an absence of evidence for genus-specific essential genes. A clear example of this sporadic essentiality is given by the set of genes regulating the σ^E^ extracytoplasmic stress response, which appears to have independently become essential multiple times in the Enterobacteriaceae. Finally, we compare our essential gene sets to the natural experiment of gene loss in obligate insect endosymbionts that have emerged from within the Enterobacteriaceae. This isolates a remarkably small set of genes absolutely required for survival, and uncovers several instances of essential stress responses masked by redundancy in free-living bacteria.

## Introduction

Gene essentiality is a core concept with importance in sub-fields spanning the breadth of microbiology and genetics (1). The motivation for the earliest attempts at determining the complete set of essential genes in bacteria, through comparisons of the first bacterial two genomes sequenced, was to determine a minimal gene set sufficient to support life (2, 3). While a somewhat esoteric object of study, the minimal gene set has important practical implications across a range of applications (4). For instance, antibiotics operate by interfering with essential cellular functions, though target-based screens for new antibiotic leads based on essential gene lists have so far met with limited success for a variety of technical reasons (5, 6). Essential gene sets are also relevant for biotechnology applications, where the essential genome puts constraints on our ability to engineer bacterial chassis (7). Finally, the construction of the first fully synthetic genomes has been deeply reliant on investigations of the essential genome (8, 9), and have been important in accelerating technological progress in synthetic biology (10).

High-throughput sequencing has accelerated the study of gene essentiality. Relatively inexpensive genome sequencing has enabled exploration of diverse reduced genomes from obligate pathogens or endosymbionts, finding bacteria that can survive with less than 200 genes, albeit under controlled conditions within host cells (11). Sequencing has also led to new experimental approaches for exploring gene essentiality, notably transposon insertion sequencing (TIS) (12) and more recently pooled CRISPR interference (CRISPRi) screens using catalytically inactive Cas proteins to silence gene transcription (13). In contrast to traditional approaches that relied on labor intensive mapping of single transposon insertion mutants (14) or targeted gene deletion (15), these approaches enable the simultaneous assay of hundreds to millions of pooled mutants in a single experiment. This has enabled the first comparisons of experimentally determined gene essentiality between strains within bacterial species (16–20), finding evidence for strain-level variation in the essential gene complement.

The Enterobacteriaceae are perhaps the single most well studied family of bacteria. They include the human gut commensal *Escherichia coli*, which has served as a model organism for 135 years (21). The *E. coli* species also includes a wide range of pathovars adapted to various intestinal and extraintestinal niches, including the polyphyletic *Shigella* lineages. Several other major human pathogens have also arisen within this family, including causative agents of typhoidal and nontyphoidal salmonellosis (*Salmonella*), dysentery (*Shigella*), yersiniosis and plague (*Yersinia*), and pneumonia and nosocomial infections (*Klebsiella*). This has led to intense interest in members of the family for medical as well as purely scientific reasons. Indeed, one of the first arrayed single gene deletion libraries (the Keio collection) was constructed in *E. coli* (15), *and has served as a major reference point for the community studying essential genes since*.

*Despite the intense interest there have been few comparisons of gene essentiality between members of the Enterobacteriaceae. Using TIS, we previously showed evidence for a conserved core of 281 genes essential in both the salmonellosis-associated Salmonella enterica* serovar Typhimurium and the typhoidal *Salmonella enterica* serovar Typhi, with a smaller set of 228 also essential in *E. coli (17)*. A similar study comparing *Shigella flexneri* with *E. coli* found a largely overlapping essential gene set, with most differences in gene essentiality being due to a loss of redundant genes or pathways in *S. flexneri* (22), *which is expected as the shigellae are simply specialized pathotypes of E. coli* (23). How these findings extend to differences in gene essentiality across the family Enterobacteriaceae is currently unclear.

To investigate gene essentiality in the family Enterobacteriaceae, we gathered a collection of previously reported and newly generated *Tn5* TIS data sets from the genera *Escherichia, Salmonella, Citrobacter*, and *Klebsiella* generated using the Transposon Directed Insertion-site Sequencing (TraDIS) technique (24, 25). We benchmark methods for identifying essential genes, developing a new classifier based on empirical clustering using DBSCAN (26). We examine how biases in TraDIS data affect gene essentiality predictions and correct for these. We compare essential gene sets between the strains in our collection, identifying a core conserved essential gene set of 201 genes and reconstructing an ancestral gene set of 296. Despite substantial variation in the essential gene content across strains, we find little evidence for private essential genes within genera. Finally, we compare our essential genes in free-living Enterobacteriaceae to related endosymbionts characterized by extreme genome reduction. We identify both a conserved core of genes absolutely required for survival, as well as a number of genes typically considered to play roles in stress responses that appear to become essential in the context of a minimal genome.

## Results

### A TraDIS gene essentiality compendium encompassing five genera of the Enterobacteriaceae

To investigate gene essentiality in the Enterobacteriaceae we assembled a collection of new and previously published data that cover five medically and scientifically important genera of the Enterobacteriaceae: *Escherichia, Salmonella, Citrobacter, Klebsiella*, and *Enterobacter* (**Figure 1**). This includes eight TraDIS libraries that have not been previously reported (see Methods), and all have been constructed in rich media with growth at 37°C. The selected *Escherichia* strains include the parent strain of the Keio collection (15), *Escherichia coli* BW25113 (27), as well as two sequence type 131 (ST131) strains associated with urinary tract infections, NCTC 13441 (28) and EC958 (29). Our *Salmonella* data includes data from the CVD908-*htrA* vaccine derivative of *Salmonella enterica* serovar Typhi Ty2 (24), the common *S*. Typhimurium lab strain SL1344 and its vaccine derivative SL3261 (17), the *S*. Enteritidis PT4 strain P125109 (33) representative of the global gastroenteritis epidemic, and S. Typhimurium D23580 (34) and A130, representatives of the two major clades involved in the HIV-associated invasive non-typhoidal salmonellosis epidemic in sub-Saharan Africa (35). We also have included *Citrobacter rodentium* ICC168 (36), a model mouse pathogen that carries the type III secretion system encoding locus of enterocyte effacement (LEE) pathogenicity island in common with the enteropathogenic (EPEC) and enterohemorrhagic (EHEC) pathovars of *E. coli* (37). *We have also included two strains of Klebsiella pneumoniae*, a major cause of multi-drug resistant nosocomial infections. These are Ecl8, a genetically manipulable model strain (38), and RH201207, a multi-drug resistant clinical isolate (39). Finally, we included a representative of the opportunistic pathogen *Enterobacter cloacae*, strain NCTC 9394.

**Figure 1.**
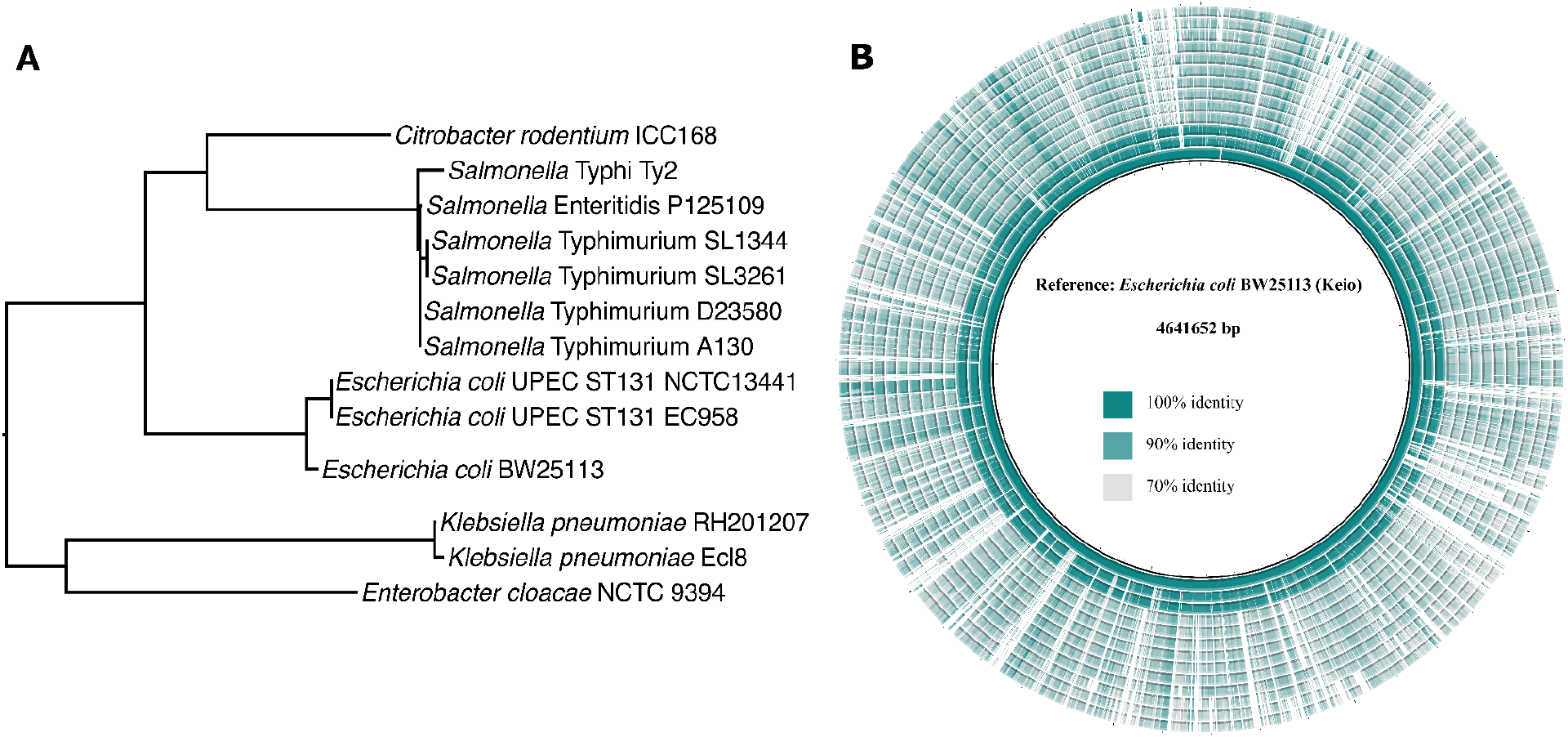
Overview of a collection of TraDIS libraries for five genera within the Enterobacteriaceae. A) An estimated tree showing phylogenetic relationships between the strains used in this study. The tree was constructed using RAxML (30) on a concatenated set of core genes (31) (see Methods). B) Genome alignment for all the genomes in this study compared to *E. coli* BW25113 from Keio collection (15), generated with BRIG (32). The genomes from the inner circle to the outer circle are *E. coli* BW25113, *E. coli* UPEC ST131 EC958, *E. coli* UPEC ST131, *S*. Typhimurium A130, *S*. Typhimurium D23580, *S*. Typhimurium SL3261, *S*.Typhimurium SL1344, *S*. Enteritidis P125109, *S*. Typhi Ty2, *C. rodentium* ICC168, *E. cloacae* NCTC 9394, *K. pneumoniae* Ecl8, and *K. pneumoniae* RH201207, respectively.

### Benchmarking essentiality classification

A number of methods have been used for evaluating the essentiality of genes using transposon insertion data. These use different features that report on gene essentiality, which can include features of the sequenced libraries themselves, such as the number of insertion sites divided by gene length (the so-called insertion index (24)), or comparisons of experimentally determined transposon read counts against synthetic reference libraries (40). Freed and colleagues benchmarked eleven of these features, finding the insertion index, mean distance between insertions, and the largest uninterrupted fraction of gene length were most predictive of essentiality (22).

Here we benchmarked four of these measures: the average distance between insertion sites in a gene, the largest uninterrupted fraction, insertion index, and the Monte Carlo method combined with DESeq proposed by Turner and colleagues (40) (**Figure 2A**). Additionally, we constructed a predictor using all four measures of gene essentiality using principal component analysis (PCA), taking the weighted sum given by the first principal component as our combined measure. To evaluate accuracy, we compared essential genes predicted from an *E. coli* K-12 BW25113 transposon library (27) with each method to curated essential genes in the EcoGene database (41). EcoGene is a refinement of essential genes determined by directed single gene knockouts in the Keio collection (15). The four measures performed similarly, as evaluated by the area under the receiver operating characteristic curve (AUC). Our PCA based predictors lead to a modest increase in performance, indicating that these measures are not completely redundant; however, given the modest improvement and the relative ease of calculation and interpretability, we have used the insertion index for further work.

**Figure 2.**
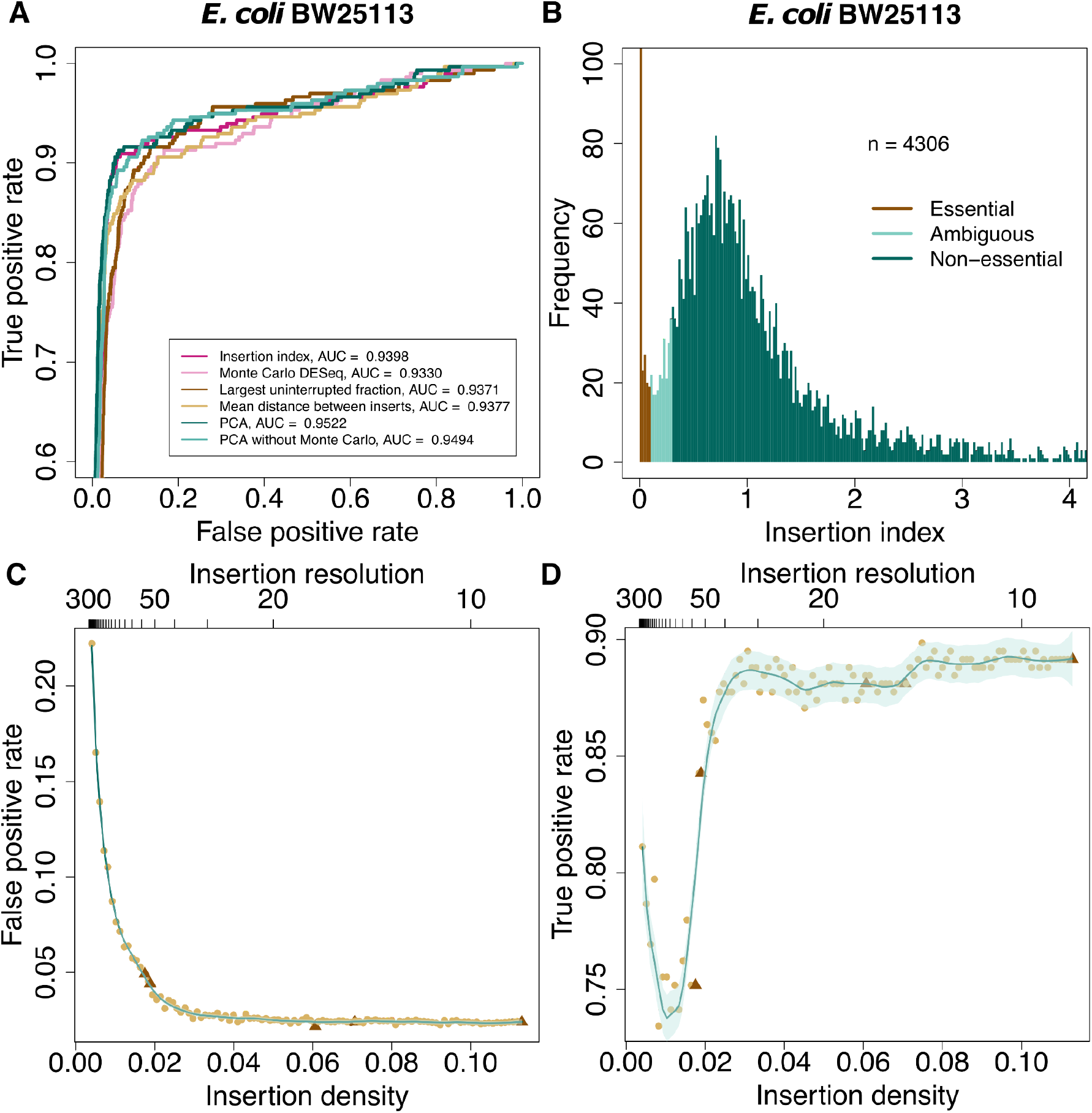
Benchmarking gene essentiality predictions. A) Receiver operating characteristic (ROC) curves showing the accuracy of 6 methods for predicting essential genes. True positives are genes that are predicted as essential for E. coli K-12 BW25113 by TraDIS and classified as essential in EcoGene. False positives are genes that are predicted as essential, but are classified as non-essential in EcoGene. Genes for which the Monte Carlo method returned NA were omitted from this comparison. B) The bimodal distribution of insertion indices illustrating essentiality classification using DBSCAN. Essential genes have the lowest insertion indices, non-essential genes have higher insertion indices, and ambiguous genes are located between these groups. C&D) Simulation of insertion density effects: The dark and light brown dots are obtained from real and down-sampled data from *S*. Typhimurium SL1344, respectively. The turquoise lines show loess curves with 0.2 span, and the light turquoise regions show the 95% confidence intervals. Insertion resolution is calculated by dividing genome length by the number of unique insertion sites. The false positive rate decreases with increasing insertion density (number of insertions divided by genome length) and remains constant after it reaches 0.04, or approximately one insertion every 25 bases (C). The true positive rate converges around an insertion density of ∼0.03, or approximately one insertion every 30 bases (D). The false positive and true positive rates are calculated by comparing predicted essential genes with the EcoGene database.

A second issue in building an essential gene classifier is choosing a threshold for essentiality. In previous work, gamma distributions have been fitted to the bimodal distribution observed in the insertion index (**Figure 2B**) and a log likelihood ratio used to determine the essentiality cut-off (24, 25). This method is heuristic, based on separation of the two component distributions and maximum likelihood fits. We have observed that this, on occasion, can lead to poor fits and instability in the classification, which can require manual intervention to correct. To make this procedure more robust, we have adopted a non-parametric clustering approach using DBSCAN (26). This leads to consistent improvements in classification accuracy compared to automated gamma fitting (25) as measured by the Matthews correlation coefficient across all our datasets using EcoGene essential genes as a gold standard (**Figure S1**).

The final component in determining gene essentiality is the transposon library itself; insufficient density will lead to genes being devoid of insertion by chance. Often, density is reported in terms of the number of insertions divided by genome length. However, this can be misleading, as insertion density is clearly non-uniform. To address this, we have constructed a series of five increasingly dense *Tn*5 libraries in *S*. Typhimurium SL1344, with an estimated 50K, 100K, 250K, 500K, and 1000K mutants based on transformation efficiency control plates. After sequencing and read-mapping, we recovered 85477, 92145, 295854, 344380, and 550657 unique insertion sites, respectively. We then computationally subsampled these insertion libraries and predicted gene essentiality using the insertion index and DBSCAN, to plot both false positive and true positive rates for a full range of library complexities (**Figures 2C&D**). These curves show that classifier performance stabilizes at around 1 transposon mutant per 35 nucleotides of genomic sequence, providing a clear minimum density for accurate prediction of gene essentiality with *Tn*5 mutagenesis.

### Evaluating and correcting for biases in *Tn*5 insertion

A number of studies have reported genomic biases in *Tn*5 insertion (16, 17, 24, 42–46). However, the findings of different studies are often contradictory, and it remains unclear whether they impact the quality of gene essentiality predictions. Using our combined data set, we examined the evidence and impact of four potential biases: positional bias within genes and genomes, and compositional bias in the form of preferred sequence motifs and G+C content.

Within genes, it has been noted that the effect of transposon insertions at the start and end of the sequence can diverge from that observed across the entire gene (14). To investigate this in our own data, we calculated local averaged insertion indices across percentiles of essential genes (**Figure 3A**). There was a clear enrichment in transposon insertions in the first 5% and last 20% of the essential gene sequences. Insertions early in essential genes may be due to either biological alternative start codons, or possibly more likely, the bias in gene prediction algorithms towards longer coding sequences (47). The enrichment of insertions at the 3′ end of essential genes may indicate that functional proteins can frequently be produced from truncated transcripts, as can be seen in the specific case of RNase E in *K. pneumoniae* RH201207 (**Figure 3B**).

**Figure 3.**
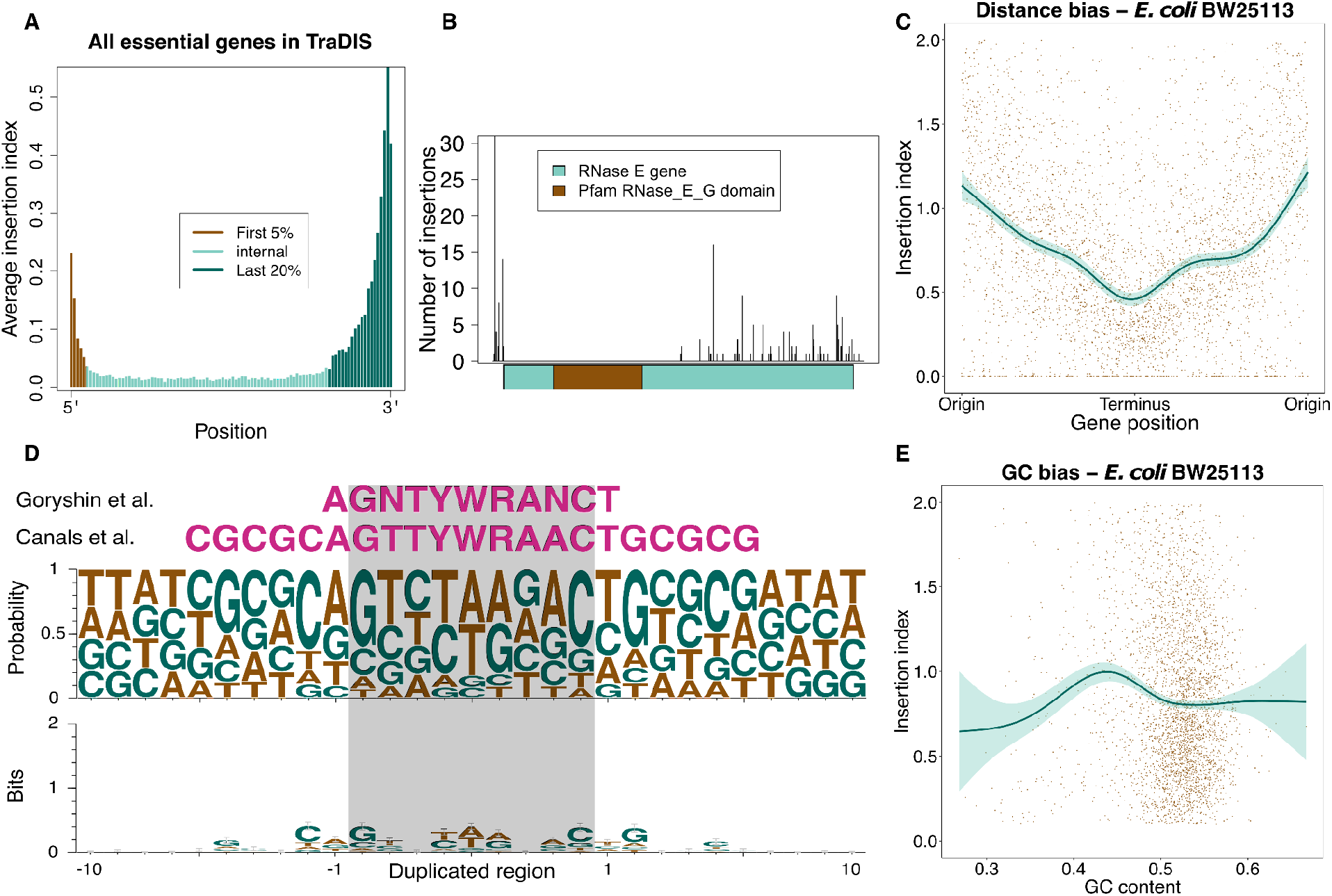
An analysis of putative sources of bias in TraDIS data. (A) The average insertion index in percentiles of all essential genes from TraDIS data. The genes are divided into 3 segments: 5% of the genes on the 5′ end (brown), 20% of the genes on the 3′ end (dark turquoise), and the rest in the middle (light turquoise). (B) Number of insertions and their location in the RNase E gene in Klebsiella pneumoniae RH201207. The 5’ end of the gene is located on the left-hand side. There are no insertions in the nuclease domain predicted by Pfam (48). (C) The insertion index versus the distance from the *dnaA* gene (usually found near the origin of replication). The turquoise curve shows the fitted GAM (Generalized Additive Model) curve, and the shading shows the 95% confidence interval. (D) Sequence logo plots generated using sequences from the 10 nucleotides flanking the 100 topmost frequent insertion sites from each genome. The relative height of each character shows their frequency (top) or the frequency of each base multiplied by the total information content of the position in bits (bottom). (E) Insertion index versus G-C content of genes. The turquoise curve shows the fitted GAM curve, and the shading shows the 95% confidence interval.

Within genomes, positional effects in transposon density have been observed with differences in density between the origin and terminus of replication (43). These differences are presumably due to the dosage effect of multiple copies of DNA near the origin of replication during exponential growth (17). We observed this phenomenon to varying degrees in our data (**Figures 3C, S2**); this variation is likely due in part to individual optimization of the transformation protocol during library creation, leading to different bacterial libraries being created in slightly different growth phases. In some genomes (e.g. *E. cloacae* NCTC9394) positional bias in transposon density did not clearly correlate with distance to the origin (**Figure S2**). This may be indicative of errors in genome scaffolding and contiguation, as the genome sequence remains unclosed.

We next examined the existence of preferred sequence motifs for *Tn*5. When the *Tn*5 transposon inserts into DNA, a region of 9 nucleotides is duplicated on each side of the transposon (42). An early study of *Tn*5 insertion based on circa 100 example insertions suggested a preferred A-GNTYWRANC-T motif (42), and this was later supported and extended by transposon sequencing in *Salmonella* (16). Other higher-throughput studies have failed to find a clear motif but have suggested a preference for elevated G+C content regions (44, 46). To address the presence of a preferred motif, we generated a sequence logo from the 10 nucleotides flanking each insertion site for the 100 most frequent insertion sites in each genome (**Figure 3D**). The nucleotide frequency observed is broadly consistent with the sequence motif proposed by Goryshin and Canals; however, the information content across the duplicated region was consistently low, suggesting this motif is not a major constraint on transposon insertion. Similarly, examining the relationship between G+C content and insertion density failed to give consistent results across genomes. Lower G+C genes tended to have slightly elevated insertion densities (**Figure 3E**), but this may be an artifact of the small number of data points below the ∼50 - 60% G+C range and the association of low G+C content with horizontally acquired DNA in enterobacterial species (27).

Correcting for the three biases most easily adjusted for (positional bias within genes, positional bias within the genome, and G+C content bias) each led to modest improvements in classifier performance. Combining the three corrections improved performance as measured by the AUC by ∼1%. However, this did lead to a substantial increase in the true positive rate in the critical low false positive range of the ROC curve (**Figure S3)**.

### Core and ancestral gene essentiality in the Enterobacteriaceae

Having investigated the factors affecting the accuracy of gene essentiality calls, we turned to investigation of essential genes in our TraDIS collection. We called essential genes in all 14 strains using the insertion index after correcting for positional bias within genes, positional bias within genomes, and G+C content bias using our DBSCAN-based classification. We then defined an ‘essentiality score’ as the log_2_ transformation of the ratio of the insertion index to the DBSCAN cut-off. Intuitively, this results in a score where values less than or equal to 0 indicate essentiality, and the value indicates 2-fold changes from this threshold. Genes with an insertion index of 0 were arbitrarily recoded to -6.5, as there were no observed scores lower than this number.

This resulted in between ∼300 and ∼440 genes classified as essential in each strain (**Figure 4A**). This is comparable to previous results in the literature applying transposon insertion sequencing to the study of gene essentiality (49), with larger essential gene sets likely reflecting the difficulty in segregating truly essential genes from those with strong growth effects in high-throughput studies. To compare essential genes between strains, we defined groups of orthologous proteins using Hieranoid (50) and a phylogenetic tree constructed using RaXML (30) from concatenated core gene alignments produced by PhyloSift (31) (see Methods). On manual examination, we found that a number of essential genes were absent in the genome sequence of *Enterobacter cloacae* NCTC 9394 due to an incomplete assembly, and this strain was excluded from further analyses.

**Figure 4.**
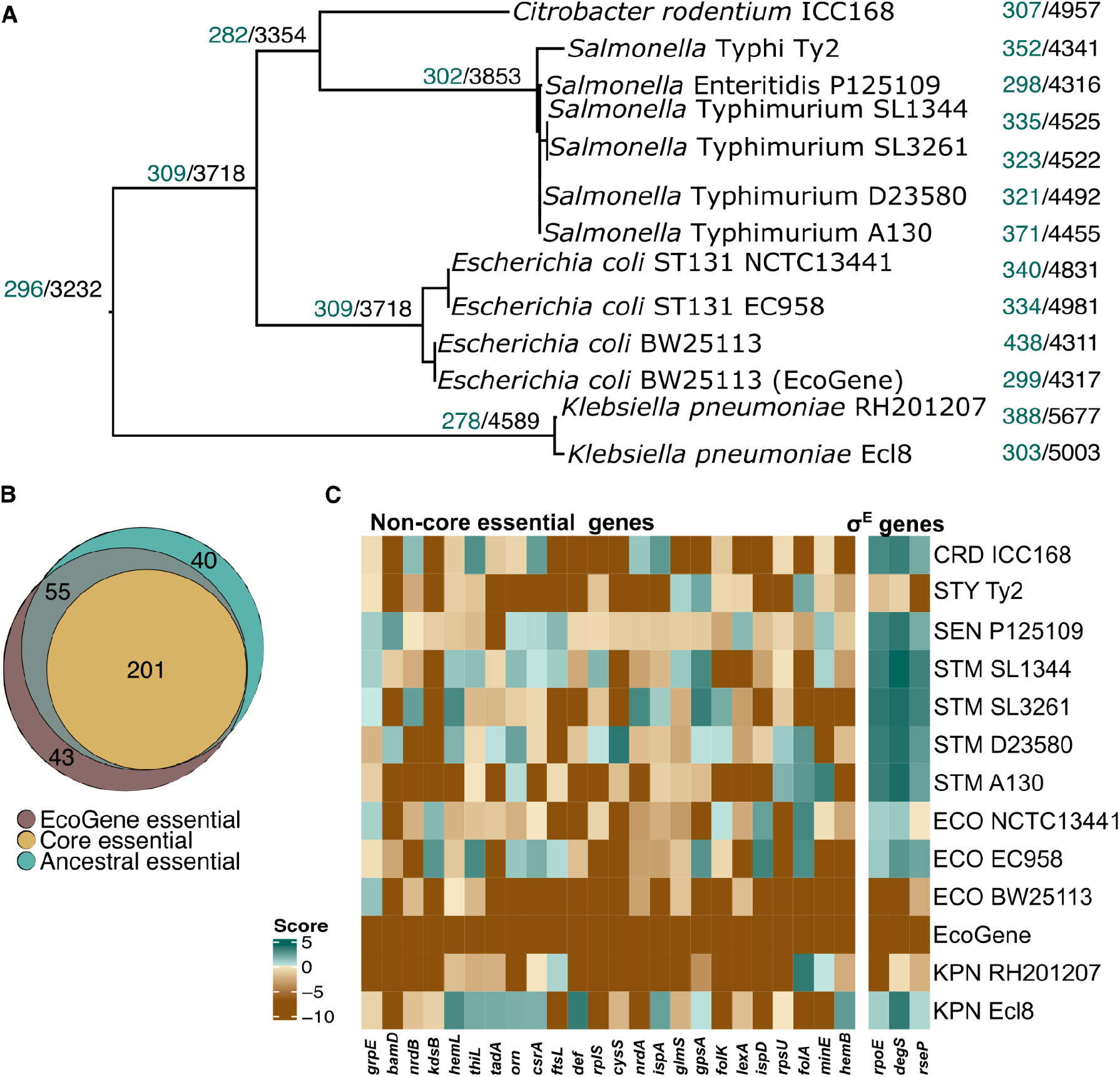
The ancestral and core essential genomes of Enterobacteriaceae. A) Reconstruction of the ancestral essential genome. Numbers on branches indicate the number of predicted essential genes (green) compared to the total number of genes predicted to be present (black) on each ancestral branch. B) Venn diagram comparing the ancestral and core essential gene sets to genes designated essential in EcoGene. C) Heatmaps illustrating variable gene essentiality across the Enterobacteriaceae. Essentiality scores are indicated by color; brown indicates essentiality while green indicates non-essentiality. Left, twenty-four genes essential in EcoGene and ancestrally essential, but not core essential, with clear evidence for non-essentiality (essentiality score > 1) in at least two strains in the collection. Right, genes involved in the σ^E^ response. CRD: *Citrobacter rodentium*; ECO: *Escherichia coli*; KPN: *Klebsiella pneumoniae*; SEN: *Salmonella* Enteritidis; STM: *Salmonella* Typhimurium; STY: *Salmonella* Typhi.

As a gold standard comparator, we included a curated collection of *E. coli* K-12 essential genes from the EcoGene database (41). Unfortunately, this resource is no longer maintained, but an archival copy is available in the Internet Archive^1^, and we additionally provide the gene list in (**Table S1**). This list curates the Keio collection of directed gene deletions (15) with observations from later studies which have investigated technical artifacts in the Keio collection, and contains 299 essential genes. Major differences from the Keio collection are the inclusion of 15 genes not considered essential by Keio which were subsequently shown to be artifacts of gene duplication, namely *glyS, ileS, alaS, coaA, coaE, dnaG, glmM, groL, parC, prfB, polA, rho, rpoD, rpsU*, and *lptB*. Genes considered essential in the original Keio paper but later shown to be non-essential include the ribosomal proteins *rpsI, rpsM*, and *rpsQ*, mutants of which grow extremely slowly (51), and *rnc* encoding RNase III, whose deletion is thought to have a polar effect on the downstream essential gene *era* with which it is co-transcribed (52).

We took multiple approaches to exploring the contents of our essential gene collection. As a first approach, we defined the core essential genes in our genomes as the intersection of essential genes across all TraDIS libraries and the EcoGene list. This resulted in a set of 201 genes essential across all strains in our collection. We also applied Fitch’s algorithm, a parsimony-based approach to reconstructing ancestral states (53), to reconstruct the set of genes essential in the ancestor of all strains included in our collection resulting in 296 ancestrally essential genes (**Figure 4A**). Core and ancestrally essential genes are summarized in Table 1.

**Table 1.**
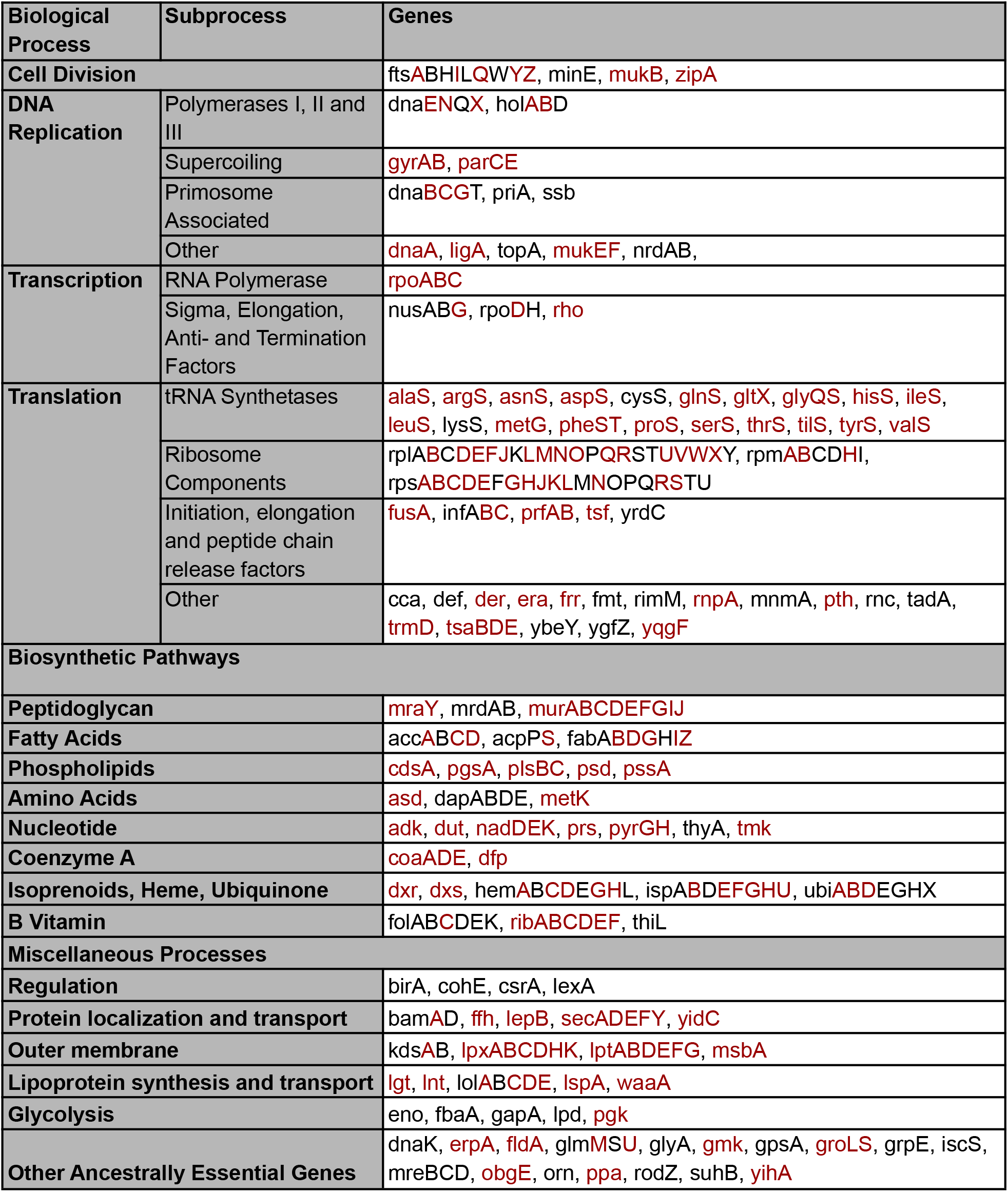
Ancestral essential gene functions in the Enterobacteriaceae. Core essential genes are highlighted in red.

We found it somewhat surprising that despite the range in the number of genes predicted to be essential across our collection, the ancestral essential gene set was of similar size to that in the EcoGene gold standard. We examined the intersection of the core, ancestral, and EcoGene gene sets in an attempt to understand the differences between them (**Figure 4B**). By design, the core essential genes are a proper subset of the ancestral and EcoGene sets, and contain ∼2/3rds of the genes essential in either of these sets. Fifty-five genes occur in both the ancestral and EcoGene sets but are not core. Manual inspection led us to conclude that two of these were excluded from the core gene set due to annotation errors: strains D23580 and A130 lacked annotations for *rpmC*, though the gene sequence appears to exist at the same locus as other *Salmonella* strains. Similarly, the A130 annotation appears to contain a truncated annotation for *accC*, which was not properly clustered by Hieranoid. The gene *cohE*, encoding a phage repressor protein, was essential and present in *E. coli* and *K. pneumoniae* strains but not others, suggesting independent horizontal acquisition of similar prophage in the two lineages and misclassification as ancestral. Twenty-eight of the remaining genes were predicted as essential in most strains, with only one or two strains scoring slightly above (< two-fold higher) the essentiality threshold, indicating that most of these genes are likely either core essential or at least cause severe fitness defects in all strains surveyed.

This analysis left twenty-four genes that were ancestrally essential and essential in EcoGene, but not classified as core essential, where at least one strain had a score greater than two-fold higher than the essentiality threshold (**Figure 4C**). Some of these had clear-cut explanations. For instance, *folA*, encoding dihydrofolate reductase, is the target of the antibiotic trimethoprim commonly used in the treatment of urinary tract infections. *folA* non-essentiality could be explained in all cases by the acquisition of alternative trimethoprim-resistant dihydrofolate reductase enzymes either in the plasmid or chromosome. Another explained example is *cysS*, encoding a cysteine-tRNA ligase uniquely non-essential in *S*. Typhimurium D23580, which was previously shown to be displaced by a plasmid-borne copy in this strain (54). Most others were less easily explained. For instance, *ispD, ispA*, and *hemL* in the isoprenoid and heme biosynthesis pathways were called as clearly non-essential in multiple strains. The gene *ispD* in particular, encoding a key cytidylyltransferase in isoprenoid biosynthesis and a target of fosmidomycin (55), was non-essential in both ST131 UPEC strains. These are particularly interesting as the methyl-D-erythritol (MEP) pathway is not present in metazoans and hence has served a target for therapeutic intervention (56).

### Evidence for differences in essentiality among the Enterobacteriaceae

Forty-three genes were essential in the EcoGene list, but are neither core nor ancestral (**Figure 4B**), constituting almost 15% of essential genes in *E. coli*. This is of particular importance, as the essential genes of *E. coli* are often taken as representative for related species. Fourteen were variably present in our strain collection, and a third of these encoded the antitoxin components of toxin-antitoxin systems (*chpS, mazE, mqsA, yafN, yefM*). Phage repressors were also in this set of essential, variably present genes as seen in previous comparisons of gene essentiality between strains (17), notably including *dicA* encoding the c2 repressor in the cryptic Qin prophage (57) that controls expression of the small protein DicB and small RNA DicF that affect cell division and phage susceptibility (58). Others include the *trpS* tryptophan tRNA-ligase, of which an ancestral second copy appears to have been lost in the lineage leading to *E. coli* and *C. rodentium*.

Approximately 60% (27/43) of the genes essential in the EcoGene list but not core or ancestrally essential were also non-essential in our *E. coli* BW25113 dataset. We cross-compared our results to an independently generated and analyzed *E. coli* BW25113 TraDIS library (59), and found that with the exception of two genes very close to the essentiality decision boundary (*yrfF* and *can*), these results agreed with ours in all cases. Some of these disagreements with EcoGene could be explained by domain essentiality within a gene otherwise tolerant of insertions on manual examination. For instance, *rne, polA, yejM*, and *spoT* all appeared to contain essential domains across multiple strains and were additionally found to be essential across a collection of *E. coli* strains in a recent CRISPRi screen (20). However the reasons behind the majority of these disagreements between EcoGene and TraDIS data remain unclear, and many of these genes remain poorly characterized.

One example of a set of genes essential in EcoGene and not core or ancestrally essential is associated with *rpoE*, which encodes the extracytoplasmic stress responsive sigma factor σ^E^ (**Figure 4C**). While σ^E^ has long been known to be essential in *E. coli* (60), *and suppressor mutants are easily obtained (61), the exact reason why it is essential remains unclear, though may have to do with the accumulation of misfolded outer membrane proteins (OMPs) in the deletion strain (62). For activation, σ*^*E*^ *requires release from the anti-sigma factor RseA via the activity of the DegS and RseP proteases. All three genes rpoE, degS*, and *rseP* were essential in our BW25113 data in agreement with EcoGene; however, none of these genes were essential in either of the ST131 *E. coli* strains included in our collection. These genes have also previously been reported to be essential in *S*. Typhi CVD908-htrA (17), confirmed both here and by an independent study in *S*. Typhi Ty2 (16). We also find all three to be essential in our clinical isolate of *K. pneumoniae* RH201207. Why this system is essential in three phylogenetically separated strains and not others is unclear. It has been speculated that σ^E^ essentiality may depend on the presence of *ydcQ/hicB* (61), *though this was later shown to be the result of HicB toxin activation (63) and neither S*. Typhi CVD908-htrA nor *K. pneumoniae* RH201207 encode the HicAB toxin-antitoxin system. The OMP content of Enterobacterial membranes can vary widely between strains, so this could possibly be a driving factor in this differential essentiality if accumulation of misfolded OMPs plays a role in differential requirement for σ^E^. Whatever the ultimate reasons for σ^E^ essentiality in these strains, it is clear that this trait has arisen independently multiple times within the Enterobacteriaceae and can occur in strains of clinical relevance.

### Analysis of essential genes conserved in reduced genomes identifies essential stress responses

Two major approaches have been taken to determining gene essentiality in the literature: experimental approaches, like the TraDIS technique used here, and comparative genomics based on the conservation of individual genes across the bacterial phylogeny. Bacteria with reduced genomes, such as obligate pathogens and endosymbionts, are particularly informative for this purpose as they can survive with just hundreds of genes (11). The family Enterobacteriaceae contains a number of obligate insect endosymbionts, including the model symbionts *Buchnera aphidicola* in aphids and *Wigglesworthia glossinidia* in the tsetse fly (64). These bacteria typically reside in specialized fat cells termed bacteriocytes, which provide nutrients to the endosymbiont in exchange for essential amino acids or vitamins absent from the insect diet. While phylogenetic inference on highly reduced genomes is challenging, it does appear the endosymbiont lifestyle has emerged independently from free-living or facultative intracellular ancestors several times within the Enterobacteriaceae (65). Investigating gene conservation in these “natural experiments” provides an orthogonal approach to our TraDIS screens for determining core gene essentiality.

We collected a set of thirty-four endosymbiont genomes from within the Enterobacteriaceae on which to conduct conservation analysis. More than half (17) were *Buchnera aphidicola* strains, but also included representatives of the *Sodalis, Wigglesworthia, Blochmannia, Baumannia* genera; two secondary symbionts of psyllids; and *Moranella endobia*, a symbiont which lives within the betaproteobacterium *Tremblaya princeps*, itself a symbiont of mealybugs (66). Using the same Hieranoid-based method as we used for our essentiality analysis, we identified orthologous genes between these strains and our collection of strains used for TraDIS analysis. This allowed us to identify genes universally conserved among endosymbionts, as well as compare them to our essential gene sets for free-living Enterobacteriaceae on rich media.

We identified a remarkably small set of 120 genes universally conserved in the endosymbionts (**Figure 5**), and 73% (88 of 120) overlapped with either our core or ancestral essential gene sets determined by TraDIS. Nearly half of these (41 genes) are ribosomal proteins, comprising all but eight of the essential ribosomal proteins in our ancestral essential gene set (**Table 1, Table S1**). Other large classes of genes essential in free-living Enterobacteriaceae and conserved in endosymbionts included a subset of aminoacyl-tRNA biogenesis (nine genes, including *trpS*), genes involved in translation (*der, frr, fmt, infA, infB, mnmA, tadA, tsaD, pth, rnc, ybeY, ygfZ*), DNA replication (*dnaB, dnaN, dnaQ, dnaX, gyrB, holB, ssb*), and fatty acid biosynthesis (*acpS, fabB, fabG, fabI*).

**Figure 5.**
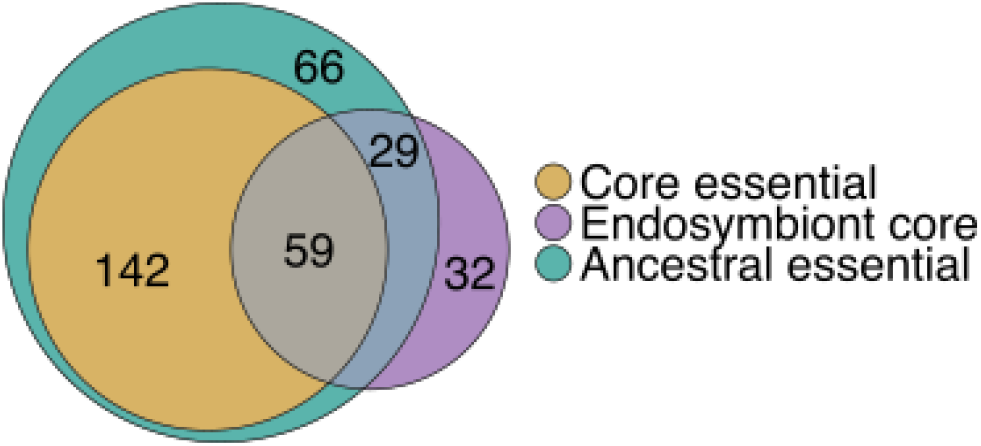
Comparing gene essentiality in free-living Enterobacteriaceae to conservation in reduced genomes. Endosymbiont core genes are defined as those universally conserved across a set of thirty-four endosymbiont genomes from within the family Enterobacteriaceae.

While endosymbiont lineages largely conserved a core of genes involved in information transfer, few essential biosynthetic pathways were universally conserved. Loss of genes involved in ubiquinone synthesis have been noted since the sequencing of the first *Buchnera* genome (67); we find that these, and other associated isoprenoid and heme biosynthesis genes involved in electron transport have been lost independently multiple times. Similarly, essential genes associated with synthesis of the outer membrane, peptidoglycan, and lipopolysaccharides have been lost multiple times, including the genes responsible for the synthesis of phospholipids. It has been speculated that the host may provide phospholipids for the symbiont membrane (68), though the details of how this may occur remain unclear. In the case of *Moranella*, it has been shown that host encoded peptidoglycan synthesis genes of diverse bacterial origin can complement the absence of these genes in the endosymbiont (69, 70).

Thirty-two genes were universally conserved in endosymbionts despite not being essential in free-living species (**Figure 5, Table 2**). Some of these could be explained by their relationship to core information transfer processes: for instance six encoding non-essential ribosomal proteins (*rplI, rpmEFGJ, rpsI*), and five encoding ribosomal- and tRNA-modifying enzymes (*gidA, mnmE, mnmG, rluD*, and *rsmH*). Others appeared to be associated with stress responses in free-living bacteria, which was unexpected as the host is generally thought to maintain a stable environment for its symbionts (68). These included for instance *lepA*, encoding a non-essential translation factor that contributes to survival of stress conditions (71) and whose deletion has previously been shown to negatively interact with deletion of ubiquinone biosynthesis genes in *E. coli* (72), *and ahpC*, encoding alkyl hydroperoxide reductase required to scavenge hydrogen peroxide in the absence of catalase (73).

**Table 2.** Genes universally conserved in endosymbiont genomes and not ancestrally essential in the Enterobacteriaceae, excluding those encoding ribosomal proteins.

| Gene | Process | Annotated protein function |
| --- | --- | --- |
| *aceF* | Glycolysis | Dihydrolipoyllysine-residue acetyltransferase component of pyruvate dehydrogenase complex |
| *ahpC* | Oxidative stress response | Alkyl hydroperoxide reductase C |
| *aroK* | Biosynthesis of amino acids | Shikimate kinase 1 |
| *clpP* | Protein degradation | ATP-dependent Clp protease proteolytic subunit |
| *clpX* | Protein degradation | ATP-dependent Clp protease ATP-binding subunit |
| *cspE* | RNA chaperone | Cold shock-like protein CspE |
| *deaD* | RNA degradation | ATP-dependent RNA helicase DeaD |
| *efp* | Translation | Elongation factor P |
| *gpmA* | Glycolysis | 2,3-bisphosphoglycerate-dependent phosphoglycerate mutase |
| *greA* | Transcription | Transcription elongation factor |
| *grxD* | Iron-sulfur cluster biogenesis | Glutaredoxin 4 |
| *lepA* | Translation | Elongation factor 4 |
| *lon* | Protein degradation | Lon protease |
| *map* | Protein maturation | Methionine aminopeptidase |
| *mnmE* | Translation | tRNA modification GTPase |
| *mnmG* | Translation | tRNA uridine 5-carboxymethylaminomethyl modification enzyme |
| *nfuA* | Iron-sulfur cluster biogenesis | Fe/S biogenesis protein |
| *prmC* | Translation | Release factor glutamine methyltransferase |
| *rluD* | Translation | Ribosomal large subunit pseudouridine synthase D |
| *rsmH* | Translation | Ribosomal RNA small subunit methyltransferase H |
| *smpB* | Translation | SsrA-binding protein |
| *sufA* | Iron-sulfur cluster biogenesis | Fe/S cluster assembly protein |
| *tkt* | Pentose phosphate pathway | Transketolase 1 |
| *tpiA* | Glycolysis | Triosephosphate isomerase |
| *trpS* | Translation | Tryptophan--tRNA ligase |
| *ycfH* | Unknown | Uncharacterized metal-dependent hydrolase |

The requirement for stress response genes extends to entire systems, including for instance the trans-translation system comprising SsrA (the transfer-messenger RNA), SmpB, and the ClpPX protease that collectively rescue stalled ribosomes (74). While generally thought to be important for the survival of stress conditions in free-living bacteria (75), tmRNA has been found in some rare mitochondria (76) suggesting it may also play an important role in reduced genomes. A final striking example is given by the *cspE*, encoding the cold shock protein CspE. As the name suggests, cold shock proteins were discovered as regulators of the cold shock response (77), and the best characterized, CspA, acts to melt RNA structures to ease translation at low temperatures (78). Free-living Enterobacteriaceae genomes encode multiple cold shock protein homologs, with *E. coli* carrying nine, for example. CspE in particular has recently been characterized as playing a key role in the virulence of *S. enterica* (79), *though the nature of this role is unclear and appears to be largely redundant with the paralogous protein CspC. A recent study of Buchnera* genomes found that *cspC* has been repeatedly independently lost while *cspE* has been maintained in different lineages (80), a finding in keeping with our expanded analysis of other endosymbiont genomes. This suggests some level of non-redundancy in CspC and E function, and further suggests cold shock proteins may collectively provide some essential function during normal growth masked by their high degree of redundancy in free-living Enterobacteriaceae.

## Discussion

An understanding of gene essentiality is critical to a wide variety of fields, from industrial biotechnology to medicine. Here we have investigated the essential genomes of thirteen strains of the Enterobacteriaceae, including the archetypal model organism *E. coli*. Our analysis found approximately one third of essential genes in any given strain changing in their essentiality status over evolution across the genera *Escherichia, Salmonella, Klebsiella*, and *Citrobacter*. Rather than being an immutable gene set, this implies a remarkable flexibility in the essential genome. Beyond this study, mounting evidence supports this view. CRISPRi screens in diverse *E. coli* strains have shown that gene essentiality varies widely between strains, and that phylogenetic distance is a poor predictor of the fitness effect of any particular gene (20); similar observation have been made using transposon mutagenesis in a collection of *Pseudomonas aeruginosa* strains (18). Even during a relatively short period of adaptation to a static environment, recent work investigating gene essentiality in the Long-Term Evolution Experiment (LTEE) found nearly two hundred genes that change in essentiality status (81). Here, we had initially expected to identify genus-specific essential genes but were unable to support this with our TraDIS dataset. While it is possible that confounding factors in TraDIS screens such as difficulties in detecting domain essentiality or protection of DNA from transposition by high molecular weight DNA-binding proteins (82, 83) may complicate the identification of genus-specific essential genes, it is difficult to explain our complete inability to find them. This suggests that the variation in gene essentiality observed between closely related strains is similar to that seen across much larger phylogenetic distances.

Despite this variation, we also found a core essential gene set comprising 201 genes whose essentiality status remained unchanged across all strains in our collection. Remarkably, similarly sized sets of conserved essential genes have been previously reported between *E. coli* and the alphaproteobacterium *Caulobacter crescentus* (84), *and even the Gram-positive Bacillus subtilis* (85). *How can we reconcile the deep conservation of this core essential gene set with the turnover observed across the other third of the essential gene set? Recent work in Streptococcus pneumoniae* investigating core and strain-dependent essential genes has suggested that the fitness context is a driving feature discriminating the two (86): while disruption of core essential genes was only ever successful through the generation of merodiploids carrying a wild-type copy, disruption of strain-dependent essential genes frequently gave rise to suppressor mutations at distal loci. This suggests the existence of more paths to non-essentiality for some essential genes than others. The variable essentiality of the σ^E^ response within our TraDIS collection is a canonical example of a system with many paths to non-essentiality, as its essentiality can be suppressed by a number of distinct spontaneous mutations (61), which may partially explain how it has transitioned between essential and non-essential states multiple times over evolution within the Enterobacteriaceae.

If fitness context defines core and accessory essential genes, then obligate endosymbionts represent an almost pathological case study. The isolation and extreme bottlenecks these organisms have been subjected to have pruned the genome of redundancy and routes for compensation, uncovering essential processes masked in the larger genomes of free-living organisms. A prime example of this is the trans-translation system for rescuing stalled ribosomes, centered on tmRNA. In *E. coli*, trans-translation is important for survival of a wide variety of stresses, including heat, acid stress, and starvation (75). Genetic interaction analysis has shown that tmRNA is only non-essential because of redundancy with other ribosome release systems (87, 88), which have been lost in the majority of the endosymbiont genomes we investigated here. The association of many genes universally conserved in endosymbionts with multiple stress responses in free-living bacteria appears to be a general trend, and suggests that at least some of what is generally considered a stress response is actually essential for cellular maintenance even in relatively static conditions, but masked by the redundancy of most bacterial genomes.

Given that essentiality is contextual, how might we best quantify this in the future? Our observations suggest several future directions. The first is investigating the relationship between conditional essentiality in response to stress, and the core essentiality of a process. Many non-essential core conserved genes in endosymbionts are known to have pleiotropic phenotypes in free-living bacteria. It seems plausible that the more unrelated phenotypic traits a gene affects, the more difficult it will be to compensate for during evolution. Indeed, genes whose deletion leads to sensitivity to a wide range of conditions tend to be involved in similar processes to essential genes (89). A second direction may be in directly reducing redundancy through genetic interaction screening. Methods for this have traditionally been extremely labor intensive, involving individually mating thousands of deletion strains (90, 91). Transposon insertion sequencing has made this analysis more straightforward for single query genes (12, 92), but would still require construction of thousands of mutant libraries for true genome-wide analysis. CRISPR array methods, which allow multiplex expression of several guides within a single cell simultaneously (93, 94), could provide a viable approach to combinatorial silencing of genes allowing for truly comprehensive screening of genetic interactions. This would allow us to peel back the layers of redundancy and come to a better understanding of the minimal requirements for cellular viability.

## Acknowledgements

LB is supported in part by a Bayresq.net project grant (Rbiotics) from the Bavarian State Ministry for Science and Art. GL and RK were supported by the BBSRC Institute Strategic Programme Microbes in the Food Chain BB/R012504/1 and its constituent projects BBS/E/F/000PR10348 and BBS/E/F/000PR10349. PPG is supported by the Marsden Fund (19-UOO-040 & 17-UOO-050), a Rutherford Discovery Fellowship (10-UOC-013), and a Ministry of Business, Innovation and Employment Grants (UOOX1709 & UOAX1932). Sequencing was supported by the Wellcome Sanger Institute and the KAUST Bioscience Core Laboratory.

## Methods

All data analysis scripts are freely available at https://github.com/Gardner-BinfLab/EnTrI. Raw sequencing reads have been deposited in GEO with accession GSE216013.

### Construction and sequencing of TraDIS libraries

All TraDIS libraries first described here were constructed using a *Tn5*-derived transposon as described previously (24). Briefly, transposons carrying either a Kanamycin (*Citrobacter rodentium* ICC168, *Enterobacter cloacae* NCTC 9394, *Klebsiella pneumoniae* Ecl8, *Salmonella* Typhimurium D23580) or Chloramphenicol (*Salmonella* Typhimurium SL1344, *Salmonella* Typhimurium A130, *Salmonella* Enteritidis P125109, *Escherichia coli* ST131) resistance cassette were incubated with EZ-*Tn5* transposase (Epicenter, Madison, USA) and electroporated into the target bacterium. Transformants were selected by plating on LB agar containing appropriate antibiotics, and harvested directly from the plates following overnight incubation at 37°C. Multiple electroporation batches were combined to produce transposon mutant libraries of high complexity.

Genomic DNA from the pooled library samples was extracted using a Qiagen Genomic-tip 100/G kit (Qiagen) and used to prepare TraDIS sequencing libraries as previously described (25). Briefly, ∼2 μg of genomic DNA was fragmented to a size of ∼300 base-pairs using a Covaris Focused ultrasonicator (Covaris), followed by end repair using the NEBNext End Repair module (NEB), A-tailing using the NEBNext dA-Tailing module (NEB), and adapter ligation using the NEBNext Quick Ligation Module with a custom splinkerette adaptor (SplA5). PCR enrichment of fragments containing transposon sequence was then carried out using transposon- and adaptor-specific primers using Kapa HiFi HotStart ReadyMix (Kapa Biosystems) for 10 to 20 cycles. PCR products were analyzed on an Agilent Bioanalyzer using a High Sensitivity DNA kit (Agilent Technologies), and by SYBR green qPCR (Kapa Biosystems). Sequencing was then performed on an Illumina HiSeq 2000, HiSeq 2500 or MiSeq using transposon-specific primers and a custom dark-cycle sequencing protocol at the Wellcome Sanger Institute or the King Abdullah University of Science and Technology Bioscience Core Laboratory (*Salmonella* Typhimurium A130 and D23580). All custom adaptors, primers, and sequencing protocols are described in (25) and publicly available at https://github.com/sanger-pathogens/Bio-Tradis/tree/master/recipes.

### TraDIS read mapping and quantification

Read mapping and quantification was performed using the Bio-TraDIS pipeline (25). FASTQ files for previously published TraDIS libraries were retrieved from the NCBI SRA (95), except for the *Escherichia coli* BW25113 data (27) which was retrieved from https://genomics.lbl.gov/supplemental/rbarseq/. The FASTQ files for non-standard multiplexed or barcoded samples (*Escherichia coli* EC958 (29), *Escherichia coli* BW25113 (27)) were modified to remove barcode sequences and leave a short transposon tag compatible with Bio-TraDIS. Both new and previously published libraries were processed in a uniform manner. Reads were trimmed and quality filtered on their right (genomic sequence) ends using PRINSEQ (Schmieder and Edwards 2011) with the command line options:

~~~
prinseq-lite.pl -fastq stdin -out_good stdout -out_bad null -min_len 30 -trim_tail_right 2 -trim_ns_right 1 -trim_qual_right 20 -trim_qual_window 3
~~~

Reads were filtered for 10 bases matching the expected transposon tag sequence, and filtered reads were then mapped against the appropriate reference genome and plasmid sequences, available at https://github.com/Gardner-BinfLab/EnTrI using the Bio-TraDIS pipeline with the following command line options:

~~~
bacteria_tradis -v --smalt_r 0 --smalt_k 8 --smalt_s 1 --smalt_y 0.9 -m 0 -f fastqs.txt -t TAAGAGACAG -r ref.fa
~~~

Insertion sites and associated read counts were tabulated in plot files appropriate for visualization with Artemis (96) and for further downstream analyses.

### Correcting for biases in the insertion index

For each gene in each organism, we calculated an insertion index as a measure of gene essentiality. The insertion index is calculated as 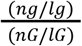 where *ng* is the number of unique insertion sites in the gene, *lg* is the length of the gene, *nG* is the number of insertion sites in the genome, and *lG* is the length of the genome. We calculated the insertion index for each gene after trimming 5% of the sequence from the 5’ end and 20% of the sequence from the 3’ end, following our analysis of insertion sites in essential genes (**Figure 3A**).

To correct for biases in the insertion index, we used fitted values from a generalized additive model (GAM). For each genome, we fitted a GAM curve (using the MGCV package in R) with the formula y ∼ s(x) to the distance vs. insertion index plot for distance normalization. The normalized insertion indices were calculated by dividing the insertion indices by the predicted insertion index from the GAM, then multiplying them by the average insertion index. The same procedure was repeated for G-C normalization.

### Classifying essential genes using DBSCAN

We used the DBSCAN R package (25) to cluster the normalized insertion indices using the DBSCAN function with parameters minPts = 200 and eps = 0.05. This density-based clustering finds two clusters for essential and non-essential genes, which are the two peaks shown by brown and dark turquoise in **Figure 2B**, respectively. A region of noise data was scattered in between these two regions shown in light turquoise. The long tail of data points on the right side of **Figure 2B** were also separated as noise by the DBSCAN algorithm, but as they have a high number of insertions, we grouped them with other non-essential genes.

### Phylogenetic tree construction

We constructed a single-copy marker gene-based phylogenetic tree for the bacteria in this study using PhyloSift (31) and RaXML (30). We first used PhyloSift-search on all genome sequences used in this study to find marker genes in PhyloSift database (31) by running `phylosift search --isolate --besthit genome_file`. Subsequently, we used PhyloSift-align to extract marker gene sequences by running the command `phylosift align --isolate --besthit genome_file`. Finally, we concatenated the alignments and used RaXMLHPC to generate a final tree by running `raxmlHPC -s protein_alignment_file -n output_tree -m PROTGAMMALG4M -p 1234 -f a -x 1234 -# 100`.

### Clustering orthologous genes

We used Hieranoid (Schreiber and Sonnhammer, 2013) with default parameters to identify orthologous genes using BLAST (97) as a similarity search tool. The RaXML species tree described previously was used for resolving orthology relationships.

### Defining ancestrally essential and ancestrally conserved genes using Fitch’s algorithm

In order to reconstruct gene essentiality along the phylogenetic tree, we used the parsimony-based Fitch’s algorithm (53) with a binary alphabet for essentiality (0 for non-essential and 1 for essential). The parent of every pair of nodes in the tree was labeled with the intersection of its children’s assigned sets, if the intersection was not empty. Otherwise, it was labeled with the union of its children’s assigned sets. We continued this process until we reached the root of the tree. At the root, if the assigned set only contained 1, the gene was considered ancestrally essential. Otherwise, it was considered ancestrally non-essential. The same procedure was performed, using gene presence rather than essentiality, to predict ancestral conservation in **Figure 4A**.

### Description of Supplementary Information

**Figure S1**: Matthew’s correlation coefficient calculated across insertion index thresholds for all 13 bacteria in this study. True positives are genes predicted as essential at a given cut-off whose orthologs are classified as essential in E. coli K-12 by the EcoGene database. False positives are genes predicted as essential at a given cut-off, but whose orthologs are not classified as essential by EcoGene. The cut-offs derived from DBSCAN clustering and fitted gamma distributions are shown in each figure. DBSCAN-derived cut-offs outperform gamma fits in all cases.

**Figure S2**: The effect of distance from the origin of replication on the insertion index. The origin of replication is assumed to be at the *dnaA* gene. Each point represents a gene and the green line shows a GAM curve fit to the data with a 95% confidence interval. Some curves suggest possible scaffolding errors (e.g. E. cloacae)

**Figure S3**: The effect of correcting various biases affecting the insertion index on the receiver operating characteristic (ROC) curve for predicting gene essentiality. As an example, normalizing for all biases increases the true positive rate from ∼0.84 to ∼0.9 at a false positive rate of ∼0.05.

**Table S1**: Complete study data, including one orthologous group as defined by Heiranoid per row, with the following columns. Keio gene name: gene name as defined in the *Escherichia coli* BW25113 genome; EcoGene Essential, Ancestral Essentiality, Core Essentiality, Conserved in Symbiont Genomes: binary data columns indicated membership of each orthologous group in the named gene set; TraDIS Essentiality columns: essentiality scores for each genome, calculated as the log_2_ ratio of the insertion index for each gene with the essentiality threshold value; Locus columns: the locus tag for each gene in each orthologous group from both TraDIS and symbiont genomes.

## Footnotes

1 https://web.archive.org/web/20170228010715/http://www.ecogene.org/?q=topic/5

