## Supplementary Figures for "High-throughput transposon mutagenesis in the family Enterobacteriaceae reveals core essential genes and rapid turnover of essentiality"

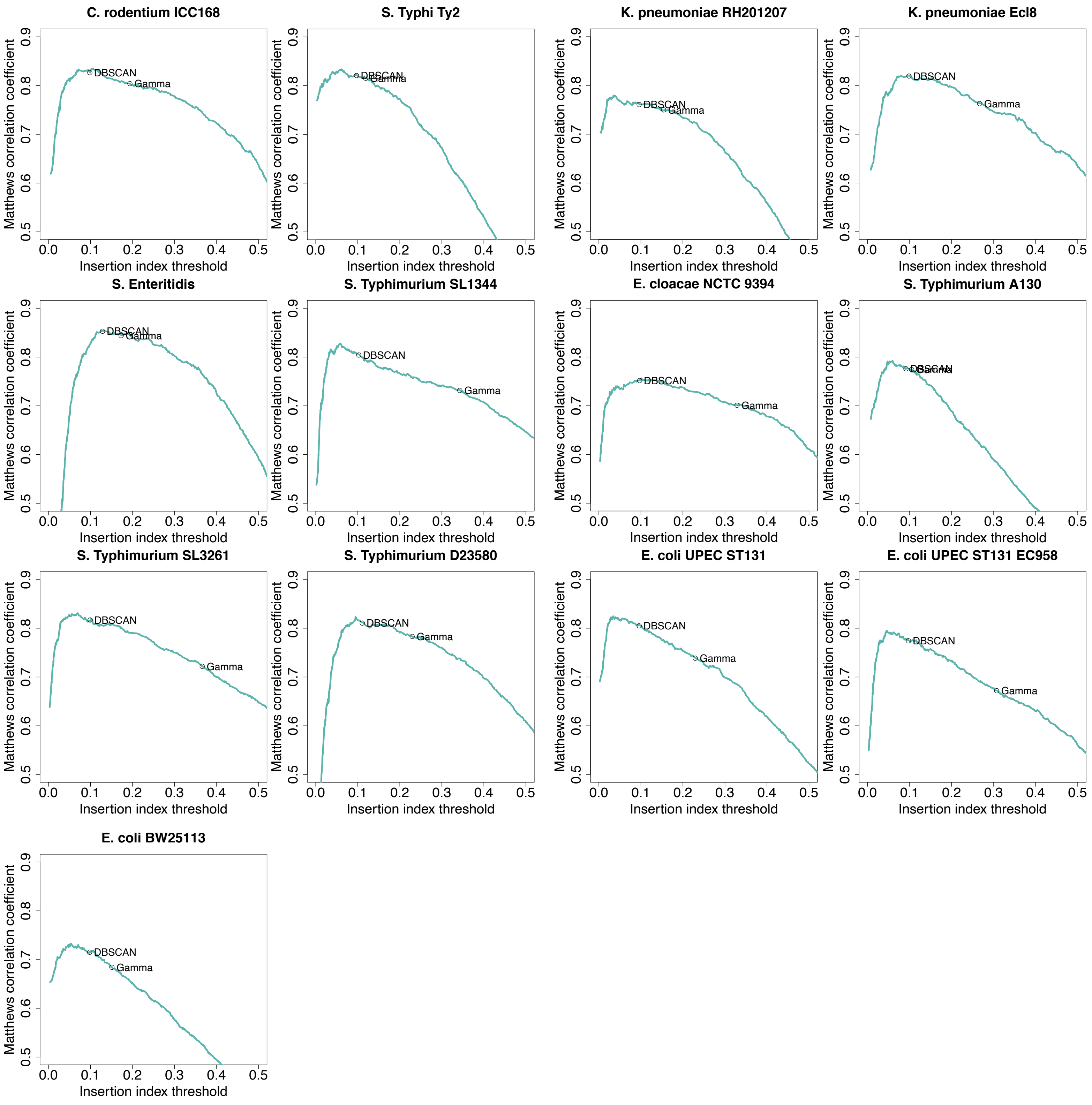


**Figure S1**: Matthew's correlation coefficient calculated across insertion index thresholds for all 13 bacteria in this study. True positives are genes predicted as essential at a given cut-off whose orthologs are classified as essential in *E. coli* K-12 by the EcoGene database. False positives are genes predicted as essential at a given cut-off, but whose orthologs are not classified as essential by EcoGene. The cut-offs derived from DBSCAN clustering and fitted gamma distributions are shown in each figure. DBSCAN-derived cut-offs outperform gamma fits in all cases.

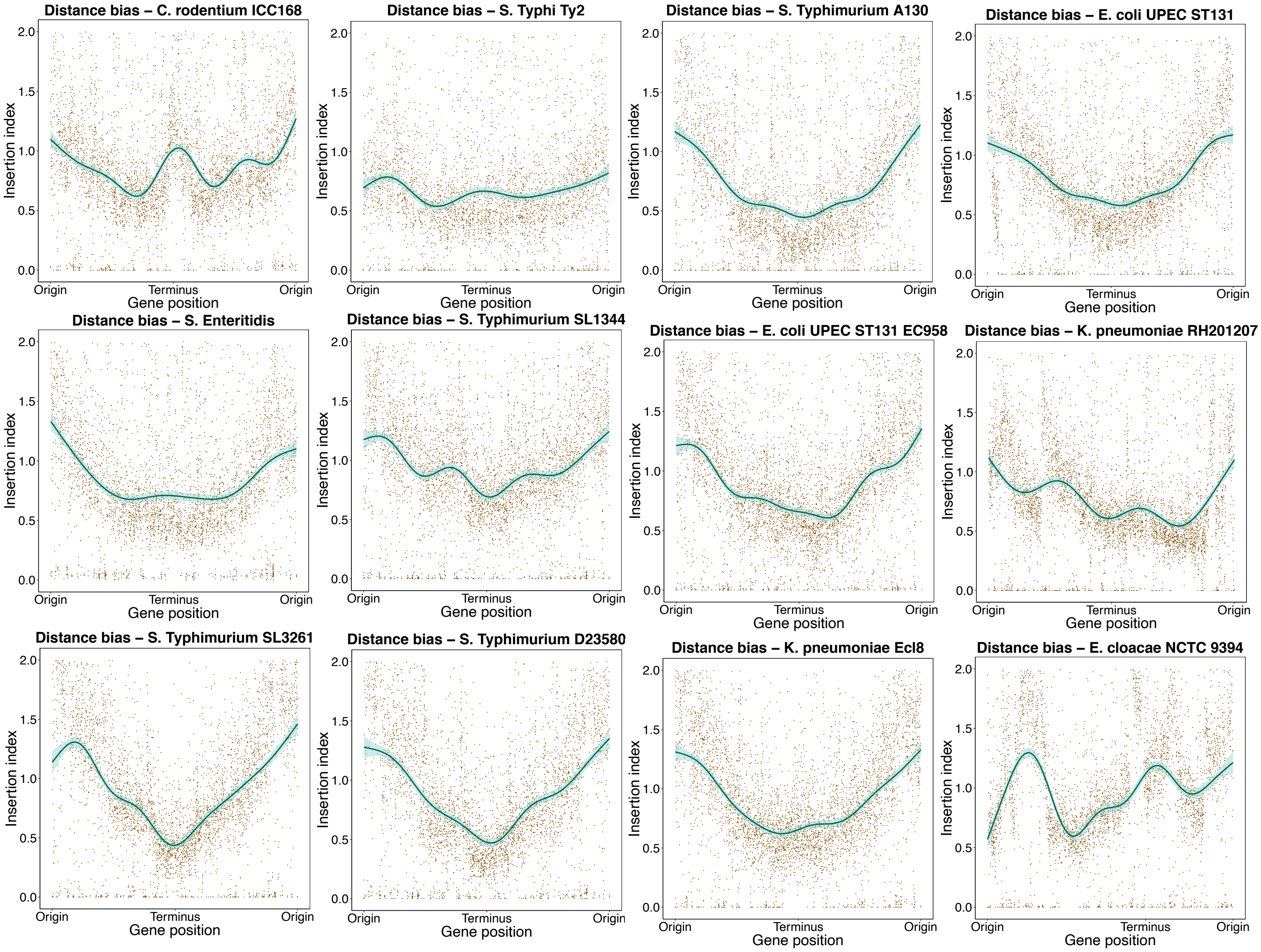


**Figure S2**: The effect of distance from the origin of replication on the insertion index. The origin of replication is assumed to be at the *dnaA* gene. Each point represents a gene and the green line shows a GAM curve fit to the data with a 95% confidence interval. Some curves suggest possible scaffolding errors (e.g. *E. cloacae*)

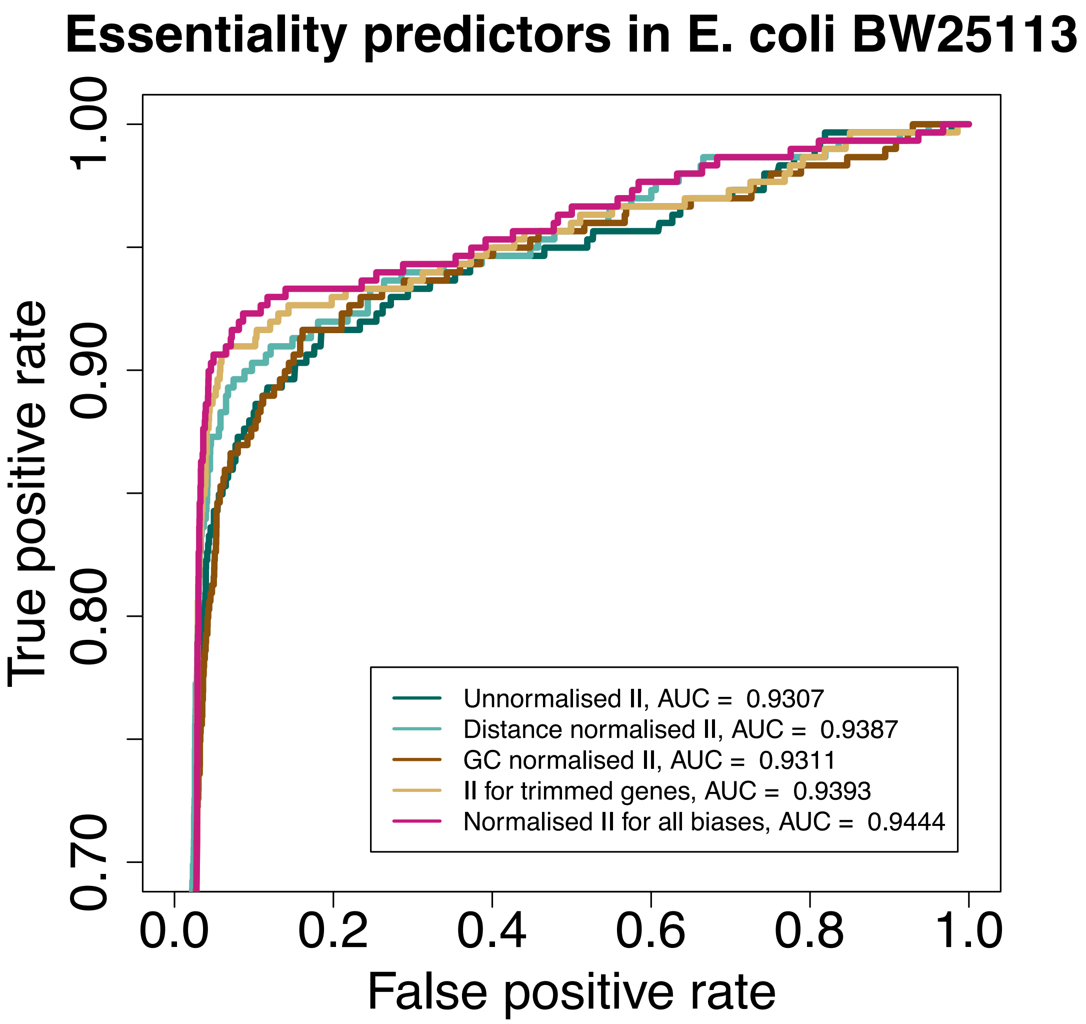


**Figure S3**: The effect of correcting various biases affecting the insertion index on the receiver operating characteristic (ROC) curve for predicting gene essentiality. As an example, normalizing for all biases increases the true positive rate from ~0.84 to ~0.9 at a false positive rate of ~0.05.
